# Common neural functions during children’s naturalistic and controlled laboratory mathematics learning

**DOI:** 10.1101/2022.01.07.475365

**Authors:** Marie Amalric, Jessica F Cantlon

## Abstract

A major goal of human neuroscience is to understand how the brain functions in the real world, and to measure neural processes under conditions that are ecologically valid. A critical step toward this goal is understanding how brain activity during naturalistic tasks that mimic the real world, relates to brain activity in more traditional laboratory tasks. In the present study, we used intersubject correlations to locate reliable stimulus-driven cerebral processes among children and adults in a naturalistic video lesson and a laboratory forced-choice task that shared the same arithmetic concept. We show that relative to a control condition with grammatical content, naturalistic and laboratory arithmetic tasks evoked overlapping activation within brain regions previously associated with math semantics. The regions of specific functional overlap between the naturalistic mathematics lesson and laboratory mathematics task included bilateral intraparietal cortex, which confirms that this region processes mathematical content independently of differences in task mode. These findings suggest that regions of the intraparietal cortex process mathematical content when children are learning about mathematics in the real world.

## Introduction

When children learn mathematics at school, they need to combine many pieces of information such as the teacher’s verbal explanation, formulas or diagrams drawn on the blackboard, reference to a book, etc. In contrast to the richness of the school environment, most of what we know about mathematics development in the brain has been established using highly controlled laboratory paradigms that tend to minimize context and extraneous input. While controlled paradigms are essential to make pure contrasts and test specific hypotheses, they can result in oversimplified and somewhat narrow theories with only limited implications to children’s real-life mathematics development (Cantlon, 2020). An alternative approach is to use visually and auditorily rich naturalistic paradigms, typically video watching, to mimic real-world learning conditions as closely as possible. This approach is essential for understanding how human brains function in the real world.

Recent neuroimaging studies have tested children with naturalistic educational videos to measure the similarities between neural processes in children and adults (Cantlon & Li, 2013; Emerson & Cantlon, 2015; Kersey et al., 2019; Lerner et al., 2021; Richardson et al., 2018). Those studies use intersubject correlation measures to quantify the similarity in brain activation timecourses between children and adults. In intersubject correlation, the neural timecourse at each voxel is tested for temporal correlation among participants. Significant correlation in brain responses among participants indicates the presence of a reliable stimulus-driven neural process in that brain region. These studies established that naturalistic paradigms evoke patterns of activation that associate or dissociate by function and age in predictable and reliable ways. For example, Cantlon and Li (2013) showed that 4-to 8-year-old children exhibited correlated brain activity - also called neural similarity - with adults while watching educational “Sesame Street” videos. Neural similarity between a given child and a group of adults is thought to indicate how adult-like is this child’s brain activity, and is thus a measure of neural maturity. In Cantlon and Li’s study, children’s neural maturity in the parietal cortex predicted their performance on mathematical tests, while their neural maturity in Broca’s area predicted their performance on verbal tests. Kersey et al. (2019) confirmed the functional dissociation between math-versus reading-related natural viewing activity, and demonstrated that both math- and reading-related networks also show immature neural activity patterns - i.e. dissimilar brain activity - in 4 to 8-year-old children relative to adults.

However, in prior naturalistic studies of children, it is unclear how brain activity during naturalistic tasks relates to activity in the more traditional simplified tasks because they were not directly compared. On the one hand, both types of tasks could recruit similar brain circuitry. For example, in 2013, Dastjerdi et al. showed that the neural populations activated during a laboratory arithmetic task, also activated when adult patients were referring to objects with numerical content during natural dialog. On the other hand, critical features of the context of learning may be unique to naturalistic video watching tasks (Cantlon, 2020). Indeed, the theory of mind network that Richardson et al. (2018) revealed with a naturalistic task differed from the one predicted by traditional tasks. Moreover, Cantlon and Li (2013) showed that math achievement was better predicted by children’s brain responses to a naturalistic math task than to a laboratory math task. Note that the two alternatives presented here are not mutually exclusive. If the same brain regions are recruited for both types of tasks, they are not necessarily recruited in the same way.

In the present study, to test whether the same brain regions that are engaged during controlled tests of a math concept in the laboratory are also involved when children are taught a lesson about that concept at school, we compare a naturalistic math video lesson and a simplified laboratory math task which both present the same arithmetic concept: the commutative principle of multiplication. In the naturalistic video, a cartoon teacher explains the formal arithmetic principle of commutativity in an educational narrative, whereas in the simplified laboratory task, the same principle is queried in a traditional two-alternative forced-choice task. Beyond the perhaps trivial difference in their respective involvement of the motor cortex versus auditory and language-related brain regions, our core prediction is that, relative to a naturalistic control video lesson testing non-mathematical content (a grammar lesson), the naturalistic and laboratory arithmetic tasks should evoke overlapping activation within brain regions that process math semantic content. These brain regions are expected to be primarily the intraparietal sulcus (IPS), as well as some frontal regions such as the inferior frontal gyrus (IFG) that are consistently found in studies of mental arithmetic and math processing in both adults and children, and the posterior inferior temporal gyrus (pITG) in adults (for meta-analyses see Arsalidou et al., 2018; Arsalidou & Taylor, 2011; Hawes et al., 2019; Houdé et al., 2010; Kaufmann et al., 2011; Yeo et al., 2017). In these regions, we then investigate similarities and differences in the activation elicited by both types of math tasks.

## Materials and Methods

### Sample size and Participants

Based on previous studies evidencing the neural dissociation between mathematics and general knowledge semantics, and others using intersubject correlations in children, we aimed at collecting data in at least 15 adults and 20 children. Data collection was interrupted due to major external events independent of the study, after scanning 39 participants: 24 typically developing children at the end of their 3^rd^ grade year (mean age 9.25 year-old ± 2 months), and 15 adult undergraduate students. One adult was excluded from further analyses due to technical issues during the fMRI exam, as well as 6 children excluded because of opting-out (4 participants) or excessive (> 3mm) head-motion (2 participants), leaving a total of 32 participants.

All adult participants and parents of child participants gave informed consent after reading or being read consent information. All children also gave their assent to participate after being read the consent information. The protocol was approved by the Carnegie Mellon Institutional Review Board.

### Protocol and stimuli

Participants were scanned using fMRI while completing a two-alternative forced-choice math task, a naturalistic mathematics video lesson, and a naturalistic grammar video lesson. While the controlled laboratory math task and math video lesson shared content about the commutative principle of multiplication, the grammar and math videos, in contrast, shared superficial perceptual features of the naturalistic video lesson.

#### Forced-choice same/different math task

The fMRI session started with 3 runs of the forced-choice task, which tested participants’ performance in an arithmetic task with the commutative principle of multiplication. Each run included 24 trials. On each trial, participants were given a few seconds to decide whether two symbolic operations or two dot arrays, including commutative pairs (e.g. “2×3” versus “3×2”), were numerically equal or not (Figure 1).

**Figure 1.**
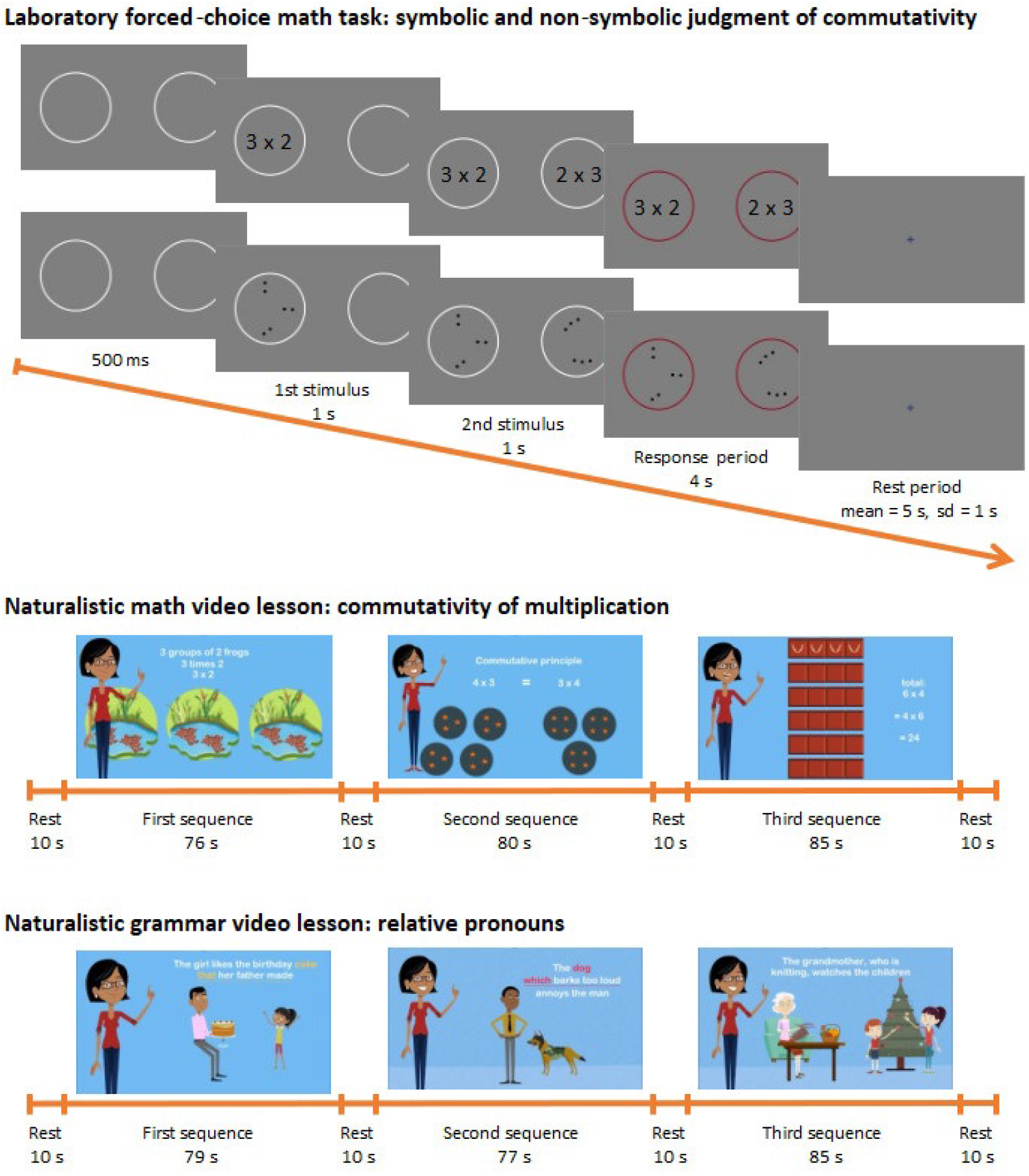
Protocol. (Top) Timeline of a trial of the two-alternative-forced-choice math task. Participants were given 4 seconds to decide whether the results of two simple operations or the numbers of dots in two different sets were the same or were different. (Middle and Bottom) Timeline of the presentation of both naturalistic arithmetic and grammar video lessons and representative frames extracted from each of the three sequences composing the videos.

Each trial of the same/different task started with two white circles horizontally distributed on a gray background, and a blue fixation cross at the center of the screen. After 500 ms, the fixation cross disappeared, and a first operation or dot array appeared in the first circle. At 1500 ms, a second operation or dot array appeared in the second circle. One second later, the two circles turned to red, indicating the beginning of the 4 seconds response period. The two operations or dot arrays remained on the screen during the entire response period. Participants were instructed to press a button with their right thumb if they thought that the two operations had the same arithmetic outcome, and to press a button with their left thumb otherwise. Each trial ended with a rest period of variable duration (mean = 5s, sd = 1s).

The same pairs were presented symbolically and non-symbolically in each run. Symbolic pairs were simple math operations (e.g., “2×3” and “3×2”). Their non-symbolic counterparts were sets of dots visually arranged in subgroups (e.g., ‘2 groups of 3 dots’ and ‘3 groups of 2 dots’). Subgroups of the same size in a set were arranged according to the same pattern. In each run, 8 pairs tested for the understanding of the commutative principle (e.g., “4×2” and “2×4”). We compared them to 8 pairs with the same number of items but different math operations (e.g., “4+2” and “3×2”). We also introduced 8 pairs controlling for numerosity and ensuring that participants did not simply estimate the number of items but computed exact additions and multiplications (half of them used the same operation: e.g., “4×3” and “4×2”; and the other half used different operations: e.g., “5+2” and “2×3”). See Table 1 for a list of stimuli.

**Table 1.** Operations used in the laboratory forced-choice math task.

| Second item | First item |  |  |  |
| --- | --- | --- | --- | --- |
|  | Same number -<br>Different number of sets |  | Different number -<br>Same number of sets |  |
|  | Same<br>operation | Different<br>operation | Same<br>operation | Different<br>operation |
| 2x3 | 3x2 | 1+2+3 | 2x4 | 5+2 |
| 3x2 | 2x3 | 4+2 | 3x4 | 3+4+5 |
| 2x4 | 4x2 | 1+2+2+3 | 2x2 | 1+3 |
| 4x2 | 2x4 | 1+3+4 | 4x3 | 2+3+3+4 |

#### Naturalistic video lessons

The arithmetic video was a lesson explaining the commutative principle of multiplication. The grammar video was a lesson explaining the principle of combining relative pronouns into phrases. Both the math and grammar videos were created with the online application “Powtoon”. They were designed to parallel a real-world school lesson by presenting formal information in a building narrative. Both lessons included the same virtual teacher explaining either a mathematical or grammatical principle over three sequences that followed a natural progression from a simple example, to a more complex example, to a general definition of the principle (Figure 1). The videos had matching visual and auditory frameworks (same teacher, same voice, same blue background, same transition animations and sound effects, same cartoon format, and a combination of symbols and pictures).

Video runs started with a plain blue screen with a central black fixation cross for 10 seconds, in order to measure the activation baseline before the video started playing. Both videos consisted of three sequences of 80 ± 4 seconds, separated by 10 seconds rest periods (Figure 1). Half of the participants watched the math video first and the other half watched the grammar video first.

### fMRI acquisition and analyses

Functional images were acquired on a Siemens Prisma 3-Tesla scanner with multiband imaging sequences (multiband factor = 4, slice acceleration factor = 3, 60 interleaved axial slices, 3 mm thickness and 3 mm in-plane resolution, TR = 2000 ms, TE = 30 ms), with 64 channel head-coil. fMRI data were preprocessed and entered into two-level analyses. At the first level, we evaluated the level of activation to each task using a standard general linear model approach on the one hand, and the synchrony across participants’ brain response to each task thanks to intersubject correlations on the other hand.

#### Preprocessing

fMRI data were processed with SPM12. Functional images were corrected for slice timing, realigned, despiked, normalized to the standard MNI brain space, resampled to a 2 mm voxel size, spatially smoothed with an isotropic Gaussian filter of 4mm FMWH, and a 100-second high-pass filter was applied.

Before completing the intersubject correlation analysis, the 6 framewise displacement parameters extracted from previous realignment were regressed out. Rest periods at the beginning of each run were then trimmed, resulting in two video runs of 276 seconds each, and three forced-choice task runs of 272 seconds each. Each run was finally z-scored (standardized to zero mean and unit variance) for each voxel. An average gray-matter mask was applied prior to performing any further analyses.

#### General linear model

Timeseries from all five runs were modeled altogether by a unique regressor obtained by convolution of the canonical SPM12 hemodynamic response function (HRF) with a rectangular kernel capturing the onset and duration of all trials. Regressors of non-interest corresponded to the six movement parameters for each run. Individual contrast images for each task (laboratory math task, naturalistic math video lesson, naturalistic grammar video lesson) relative to rest were then calculated and smoothed with an isotropic Gaussian filter of 5 mm FWHM.

#### Intersubject correlations

The technique of intersubject correlation (ISC) is a data-driven analysis technique which consists in identifying brain regions where the response to a stimulus is systematic over time, i.e. correlated among participants (Nastase et al., 2019). Both within groups (children and adults) and between children and adults, the intersubject-correlation was calculated using the leave-one-out approach. This means that, at each voxel, each subject’s timeseries (i.e., each run) was correlated with the average timeseries of all other participants. This approach directly gave us five R-maps for each subject (one for the math video, one for the grammar video, and three for the laboratory math task). Correlating each child’s timeseries with the average timeseries of the group of adults gave us five additional R-maps per child, that are typically interpreted and referred to as the child’s neural maturity maps (Cantlon & Li, 2013; Kersey et al., 2019). All R-maps were converted into z-maps using the Fischer transformation. The three z-maps calculated from each run of the laboratory math task were finally averaged into one mean z-map for further analysis.

#### Group level analyses

At the group level, to evaluate similarities and differences in activation level or intersubject correlations between tasks, we performed Bayesian tests on the individual contrast maps or z-maps obtained for each task. We used the default prior that consists in a Cauchy distribution centered on 0 with a scale parameter of 0.707 (Rouder et al., 2009). This analysis gave us the Bayes Factor (BF) which indicates how many times more likely the data are under the alternative hypothesis compared to the null hypothesis. If the Bayes Factor is equal to 1, both the null and alternative hypotheses are equally likely. A Bayes Factor of BF = 1-3 (resp .33-1) provides anecdotal evidence in favor of the alternative (resp null) hypothesis; BF = 3-10 (resp .10-.33) provides moderate evidence in favor of the alternative (resp null) hypothesis, BF = 10-30 (resp .03-0.10) provides strong evidence in favor of the alternative (resp null) hypothesis (Jeffreys, 1961). Here, we only report at least moderate evidence. Bayesian statistics are robust to variation in sample size -- a Bayes Factor greater than 3 indicates that the data are reliable. In cases without enough power to detect an effect, the Bayes Factor will be close to 1 which indicates inadequate evidence in favor of either hypotheses. Thus, the Bayesian approach of model comparison presents many advantages over classical frequentist null hypothesis testing methods: it not only provides a metric of evidence both for and against the null hypothesis, it is also robust to variation in sample size (Dienes, 2014; Schönbrodt, F.D. et al., 2017; Williams et al., 2017).

#### ROI analysis

To evaluate fine-grained similarities and differences between both the naturalistic and laboratory math tasks, we conducted a series of analyses within regions where they were found to functionally overlap, i.e. in parietal, inferior frontal and inferior temporal regions. These regions are similar to regions consistently found in studies of calculation and math semantics processing (e.g., Amalric, 2021; Ansari, 2016; Arsalidou & Taylor, 2011; Hawes et al., 2019; Yeo et al., 2017). Based on these studies, we selected four bilateral *a priori* defined math-related regions of the brain: both anterior and posterior intraparietal sulcus (aIPS and pIPS), the posterior inferior temporal gyrus (pITG), and the inferior frontal gyrus (IFG). These regions were defined functionally from two independent studies, as the intersection of the contrast of ‘calculation versus sentences’ from localizer scans performed in a cohort of 83 participants (Pinel et al., 2007) - that also served to define the ROIs used by Amalric and Dehaene (2016, 2019) - and the contrast of ‘known math versus control nonmath statements’ in a cohort of 21 participants (Amalric et al., in prep). To ensure the generalizability of math-related character of these regions of interest, we verified that they included, and were even centered around the main parietal, frontal, and inferior temporal coordinates found by Arsalidou & Taylor (2011) and by Yeo et al. (2017) in their meta-analyses of calculation and number tasks.

Within each region and for each task, we extracted the intersubject correlation values (i.e., z-values) for each adult (relative to the other adults), and for each child (relative to the other children), and applied two types of analyses: direct comparisons of mean z-values using Bayesian tests, and representational similarity analysis. Representational similarity analysis is a kind of multi-voxel pattern analysis that measures the spatial similarity of the BOLD signal elicited by various stimuli at the voxel level. Here, we applied it to test the spatial similarity of the synchronous neural activity elicited by our three tasks. To do so, we evaluated for each participant the correlation of ISC values across voxels during each of our three tasks for each math-related ROI. We then used t-tests to compare the correlation values of both math tasks (naturalistic vs controlled laboratory math) versus the correlation values of both video lessons (naturalistic arithmetic vs grammar videos). A Bonferroni correction for the number of ROIs was applied to all test results.

## Results

### Behavioral results in the laboratory forced-choice math task

Children correctly answered 89.0 ± 1.84% of the trials on the forced-choice math task. Children’s accuracy did not significantly differ from adults’ accuracy (92.4 ± 2.69%, t(30) = 1.11, p = 0.275), although they generally answered slower than adults (children: 1.97 ± 0.077s; adults: 1.08 ± 0.108s; t(30) = 7.03, p < 0.001). Children failed to respond on 13.7% of the trials while adults only missed 1.98% of trials. When counting missed trials as errors, children reached an overall accuracy of 76.9 ± 2.43%, significantly lower than adults’ accuracy (90.8 ± 2.98%, t(30) = 3.58, p < 0.002). Whether considering missed trials or not, children performed significantly better than chance for each condition (symbolic/non-symbolic x 3 categories of pairs; all p*s* < 0.05), and so did adults (all p*s* < 0.001).

### Whole-brain differences in activation to the naturalistic math video lesson compared to the controlled laboratory math task and the naturalistic grammar video lesson

We first searched for activation differences between both types of math tasks on the one hand, and between both naturalistic video lessons on the other hand. Differences in both the global intensity and the intersubject synchrony of activation were evaluated. In children and adults respectively, we computed Bayes Factors (BF) to quantify the strength of evidence in favor of activation differences between the tasks. The resulting maps, thresholded at BF > 3 to indicate at least moderate evidence, are displayed on Figure 2, and the main peak coordinates are reported in Appendices 1 and 2.

**Figure 2.**
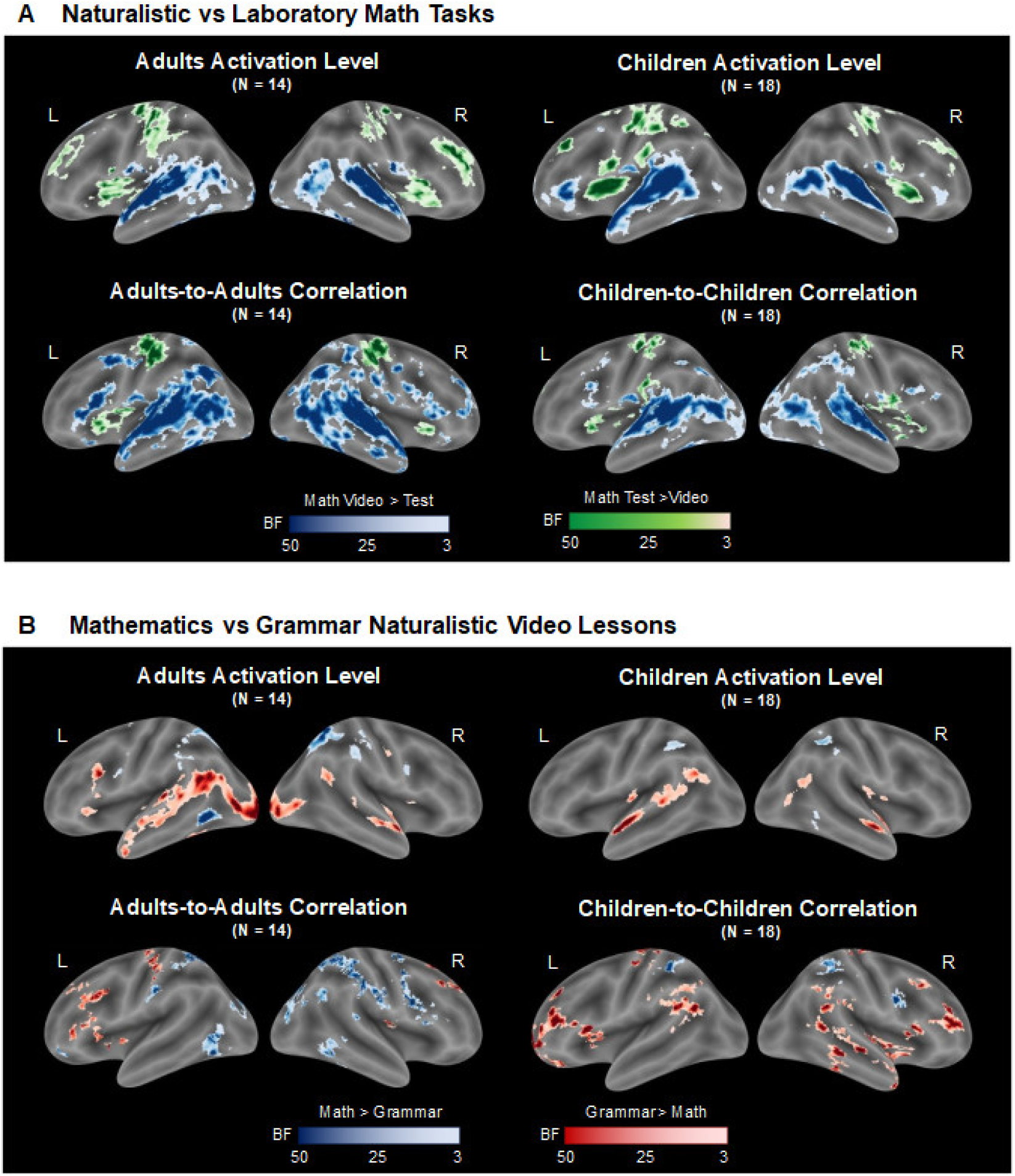
Main contrasts between tasks. (A) Whole-brain maps showing the comparison of activity elicited by both math tasks, the naturalistic video lesson in blue and controlled laboratory task in green. The top row displays comparison in activation levels obtained from a general linear model, and the bottom row displays comparison in temporally correlated activation across participants. Comparisons among adults and children are respectively displayed on the left and on the right. The maps are thresholded such that the Bayes Factor is greater than 3, showing at least moderate evidence in favor of a difference between both math tasks. (B) Same as (A) for the contrast of both naturalistic video lessons, the arithmetic lesson in blue and the grammar lesson in red.

In adults and children alike, the naturalistic math video lesson elicited greater activation than the laboratory controlled math task along the bilateral superior temporal sulcus, extending to the posterior middle temporal gyrus and to the temporal pole, as well as at various sites of the mesial superior and left middle frontal regions. In turn, the laboratory forced-choice task elicited greater activation than the math video lesson bilaterally in the motor cortex, the insula, along the brain midline in the anterior Cingulate, and along a large swath across the superior and middle frontal gyri. There was no more than weak evidence (1< BF < 3) for a difference between adults and children.

When evaluating intersubject correlations (ISC) instead of a standard general linear model (GLM), we found overall similar results, but yet noticed variations. If the swath of superior and middle frontal gyri that exhibited greater activation to the controlled math task than to the math video lesson did not show more synchronized activation, differences between the math video lesson and the controlled math task were found in a larger network when using an ISC-based approach compared to a GLM-based approach. Indeed, in addition to temporal activity, the math video lesson elicited more correlated activity than the forced-choice task in the bilateral inferior and middle frontal gyri, precuneus, and in children’s intraparietal sulcus, especially in the right hemisphere. Differences between children and adults were also found, the adults exhibiting additional correlated activity in response to the arithmetic video lesson compared to the controlled laboratory math task in the bilateral angular gyrus and the posterior inferior temporal gyrus.

We then evaluated brain regions that showed greater activity during the arithmetic video lesson than during the grammar one. For both children and adults, the resulting activation maps revealed clusters in parietal areas of both hemispheres. In the posterior inferior temporal gyrus, a large cluster was found in the adults’ left hemisphere, and a small cluster in children’s right hemisphere. Correlated activity was in parietal areas and in the right inferior frontal gyrus of children and adults alike. Additional correlated activity was found among adults in the occipital cortex, the cuneus, and the bilateral posterior inferior temporal gyri.

The converse contrast of grammar versus arithmetic video lessons revealed greater variations between the GLM-based and ISC-based approaches. The grammar lesson induced more activation than the math lesson for both children and adults along the left superior temporal sulcus, in the left inferior frontal gyrus, and in the right posterior superior temporal sulcus and temporal pole. Activation of the left superior temporal sulcus however did not exhibit stronger intersubject correlations. Grammar-related correlated activity was found in the postcentral gyrus and in various frontal regions for both children and adults. In adults, such frontal clusters were mostly in the left inferior frontal gyrus pars triangularis, and in the superior frontal gyrus. Children exhibited extensive bilateral grammar-related correlated activity in the middle and inferior frontal gyri, at the temporo-parietal junction, and along the right superior temporal sulcus.

If both GLM- and ISC-based approaches yielded comparable results in various regions of the brain, they also exhibited clear differences. While traditional GLM analyses are good for identifying differences in the amplitude of activation between conditions, they cannot capture differences in the fluctuation of neural activity over time. On the contrary, intersubject correlation statistics have the potential to reveal regions showing common patterns of activity over time across participants. Here, especially in math-related regions of the brain, the ISC-based approach is more sensitive than GLM-based analyses. In the following, we use the ISC-based approach.

### Functional overlap between the naturalistic and the controlled laboratory mathematics tasks in relation to the naturalistic grammar task

Next, we evaluated the functional overlap between our three tasks, with the ultimate goal to compare neural responses elicited by the laboratory forced-choice arithmetic task to those from the naturalistic arithmetic video lesson. ISC commonalities between both math tasks are represented in light blue on Figure 3. They are distinguished from the general overlap of all three tasks that shared a visual input modality (light green areas on Figure 3), and from the overlap of both videos that had auditory inputs (pink areas on Figure 3).

**Figure 3.**
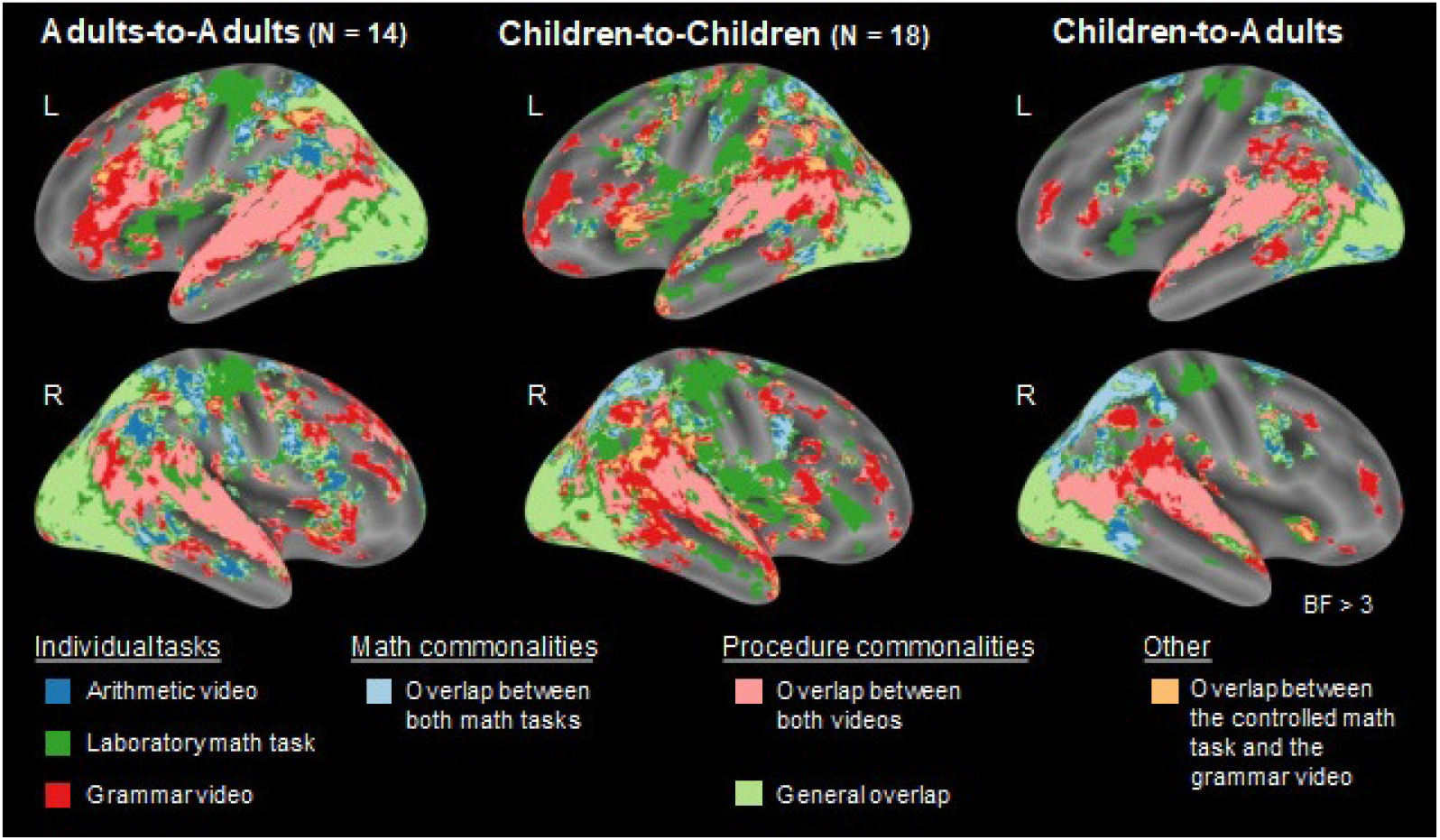
Functional overlap of all three tasks. Maps showing the significant correlations of brain activation timecourse during each task, and their respective overlaps. Red, dark blue and dark green clusters reveal correlated activation respectively associated with the grammar video lesson, the arithmetic video lesson, and the laboratory forced-choice math task. Light blue clusters, mostly found in the parietal cortex, represent correlated activation common to both math tasks, while pink clusters, mostly found in the superior temporal cortex, represent correlated activation common to the viewing of both educational videos. Light green clusters locate the brain regions exhibiting significantly correlated activation for all three tasks together. All maps are thresholded at BF > 3.

All three tasks overlapped in the occipital cortex, fusiform gyri, precuneus and inferior parietal lobule in both adults and children. In adults, this general overlap also extended to the left inferior frontal gyrus pars opercularis. Correlated activity between children and adults for all three tasks minimally overlapped in the inferior and middle occipital cortex, the fusiform gyri, and in two bilateral small sites of the posterior inferior parietal lobule.

The arithmetic and grammar video lessons overlapped in the lingual gyri, bilateral superior and middle temporal gyri, and bilateral inferior frontal sites (particularly in the left hemisphere in adults). Children’s adults-like neural responses that were common to both arithmetic and grammar video lessons were also found mostly in the lingual gyri, bilateral superior and middle temporal gyri, and bilateral inferior frontal sites. This suggests that visual and auditory processing of the videos happened similarly in children and adults.

We finally observed significant overlap of correlated activity for the naturalistic and laboratory math tasks among children in the bilateral parietal cortex, and the right inferior frontal gyrus. Adults showed the same functional overlap patterns as children but showed additional overlapping activation between tasks in the right posterior inferior temporal gyrus. Both types of math tasks elicited adult-like neural responses in children’s bilateral intraparietal sulcus, especially in the right hemisphere, in the left inferior frontal extending towards the precentral gyrus, and in the right posterior inferior temporal gyrus.

### Similarities and differences in neural responses elicited by the naturalistic versus the controlled laboratory mathematics tasks in math-related regions of the brain

The spatial overlap of neural responses elicited by both naturalistic and laboratory math tasks indicated the common involvement of brain regions similar to regions known to be generally involved in math-related processes. Here we analyzed neural response patterns within those regions to compare spatial patterns of activation. Within eight main math-related regions of interest (bilateral aIPS, bilateral pIPS, bilateral pITG, and bilateral IFG), we first measured the similarity between spatial patterns of ISC activity across tasks. For each subject, across all voxels of our math-related ROIs, we computed the correlation coefficients between the ISC values observed during each pedagogical video as well as during the forced-choice math task. We then compared the correlation of ISC values corresponding to the two math tasks versus the correlation of ISC values corresponding to the two videos (paired t-tests). In children and adults, intersubject correlations in the left and right IPS were more spatially correlated between the naturalistic arithmetic video lesson and the laboratory forced-choice arithmetic task than between the naturalistic arithmetic and grammar video lessons (Left aIPS: t(31) = 3.46, uncorrected p = 0.0016, Bonferroni corrected p = 0.0127; Right aIPS: t(31) = 2.37, uncorrected p = 0.0241, Bonferroni corrected p = 0.193; Right pIPS: t(31) = 6.39, uncorrected p < 5.10^−7^, Bonferroni corrected p < 5.10^−6^). These relations hold for children and adults separately (p*s* < 0.03).

Next, we evaluated the similarities and differences in the synchrony of activation elicited by the two math tasks within each math-related region of interest (see Figure 4). We computed the Bayes Factor (BF) of a comparison between mean ISC values from the naturalistic and controlled laboratory math tasks to quantify the strength of evidence in favor of the null hypothesis, “naturalistic and controlled math tasks elicit similar correlated activity”, versus the alternative hypothesis, “naturalistic math task elicits more correlated activity than the controlled math task”. In both adults and children, we found that the naturalistic and controlled laboratory math tasks elicited similar correlated activity in the right posterior IPS (Adults: BF = 0.271; Children: BF = 0.252). In children, they elicited similar correlated activity in the left posterior IPS (BF = 0.256), and a difference in activity in the right anterior IPS (BF = 5.75). In adults, both types of math tasks elicited similar correlated activity in the left anterior IPS (BF = 0.277), and the left IFG (BF = 0.273), while the naturalistic math video lesson elicited more correlated activity than the controlled laboratory math task in the right IFG (BF = 8.18), and bilateral pITG (Left: BF = 21.3; Right: BF = 110). This difference in the bilateral pITG was greater in adults than in children (Left: BF = 5.09; Right: BF = 5.56). We note that no such differences were found when comparing correlated activation induced by the grammar video lesson versus the controlled math task. This allowed us to verify that the difference we observed between the naturalistic and the controlled laboratory math tasks was not directly explained by video features unrelated to the content, such as animacy, color etc. Distilling across these findings, the naturalistic mathematics lesson elicited a broader pattern of systematic activation in both children and adults than the controlled laboratory mathematics task.

**Figure 4.**
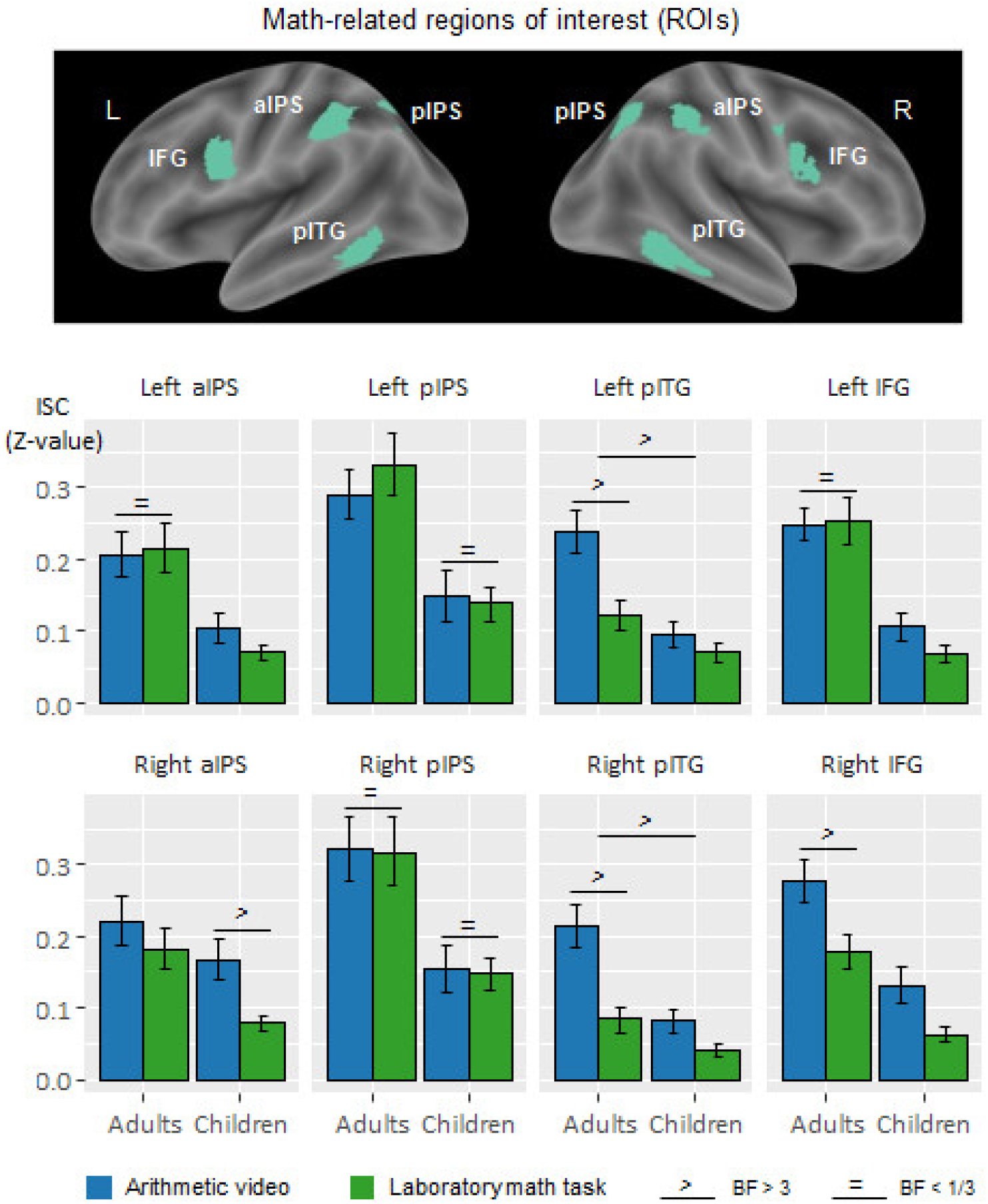
Similarities and differences in the systematic response elicited by the naturalistic math task compared to the controlled math task in math-related regions of interest. (Top) A priori defined math-related regions of interest composed of the left and right anterior intraparietal sulcus (aIPS), left and right posterior intraparietal sulcus (pIPS), left and right posterior inferior temporal gyrus (pITG), and left and right inferior frontal gyrus (IFG). (Bottom) Mean intersubject correlation values (z-values) extracted from each math-related region of interest, for the naturalistic arithmetic video lesson in blue and the controlled laboratory math task in green. Error bars equal one standard error of the mean. Equals and greater signs indicate at least moderate evidence respectively in favor of an absence of difference and in favor of a difference, between tasks as well as between adults and children.

## Discussion

We compared children’s neural activity during a naturalistic arithmetic video lesson, in which information unfolds in a rich narrative, to their neural activity during a more traditional laboratory two-alternative-forced-choice task. Intersubject correlation measures were more sensitive to detect math-related brain activity than a general linear model of the level of activation. Intersubject correlation measures revealed reliable neural signals across participants in the parietal cortex during the two types of mathematics tasks, and this pattern differed from that of the grammar task. Within the math-related network, the naturalistic math video lesson elicited more widespread systematic activation than the controlled laboratory task, especially in the right anterior IPS among children.

Regions of the parietal cortex, particularly the IPS, are often implicated in arithmetic processing (Ansari, 2008; Arsalidou & Taylor, 2011; Dehaene, 2004; Peters & De Smedt, 2018) and more generally in conceptual processing of mathematics (Amalric & Dehaene, 2016, 2019; Cantlon & Li, 2013; Vogel et al., 2015; Zhang et al., 2012). Mathematical expertise and giftedness are supported by enhanced connectivity and generally greater activity of the posterior parietal cortex (Jeon et al., 2019; O'Boyle et al., 2005; Prescott et al., 2010). Arithmetic training impacts the structural and functional properties of the IPS in children (e.g., Jolles et al., 2016; Rivera et al., 2005; Zamarian et al., 2009) and weak IPS responses during arithmetic problem-solving are associated with developmental dyscalculia (Ashkenazi et al., 2012; Price et al., 2007).

Here we confirm and extend those prior findings by comparing two tasks in which the semantic content is the same, but the format of the stimuli differs – both tasks target the same arithmetic principle of commutativity, but one is a naturalistic educational lesson and the other a forced choice laboratory task. Both the naturalistic and laboratory mathematics tasks systematically modulated neural activation in the IPS among participants, and dissociated from the grammar task. Findings from our ROI analysis suggest that both types of math tasks engaged the IPS in a systematic way over time across participants, and evoked spatially similar patterns of activation. Previous studies found functional overlap in IPS regions for symbolic and nonsymbolic numerical processing in children (Ansari, 2008; Cantlon, 2015; De Smedt et al., 2013; Holloway et al., 2010), and neural responses during arithmetic processing in children dissociate from control tasks in those regions (Bugden et al., 2012; Kersey et al., 2019; G. R. Price et al., 2013). In adults, the cortical networks for high-level mathematics processing and basic numerical processing functionally overlap, and dissociate from networks involved in language processing (Amalric & Dehaene, 2016, 2018; Baldo & Dronkers, 2007; Kanjlia et al., 2016; Kersey et al., 2019; Klessinger et al., 2007; Maruyama et al., 2012; Monti et al., 2012). The similarities in the underlying neural processes across a range of mathematics tasks from many studies are argued to be semantic in nature (e.g., Ansari, 2008; Dehaene & Cohen, 2007). Similarly, in the current study, the functional overlap and similarity between the naturalistic and controlled mathematics task, but not the grammar task, likely reflects the shared arithmetic content between the mathematics tasks.

The naturalistic and laboratory tasks exhibited some differences in various regions of the brain. The controlled laboratory task elicited greater activation than the naturalistic math video lesson in the motor cortex while the math lesson elicited greater activation in the auditory and language-related brain areas. These differences likely reflect the difference in modalities between both math tasks: participants answered by pressing buttons in the controlled math task, and they followed a verbal narrative in the math video lesson. Activation to the controlled forced-choice task was also found in the bilateral insula, and across the superior and middle frontal gyrus. These regions are often found in meta-analyses of math activity, and they have respectively been associated with participant’s intrinsic motivation in the task, and active planning and working memory processes (Arsalidou et al., 2018; Delazer et al., 2003; Houdé et al., 2010; Simon et al., 2002; Uddin & Menon, 2009). Differences between both math tasks in the insula remained when evaluating the level of synchrony in activation among participants, but disappeared in the superior frontal gyrus. This might suggest that while the laboratory math tasks required more planning than the naturalistic math video lesson, temporal fluctuations of planning-related activity were disparate among participants. Overall, the differences we observed between both types of math tasks are likely due to general modality differences and to the active character of the forced-choice task compared to the passivity of natural viewing.

Children and adults showed largely similar patterns of synchronized activation during both the naturalistic and controlled mathematics tasks within the math-responsive brain network. Both groups systematically engaged the IPS and inferior frontal regions for both the naturalistic and laboratory controlled mathematics tasks. However, unlike in children, the naturalistic mathematics task yielded greater activation in adults than the controlled mathematics task in the posterior inferior temporal gyrus (pITG). In adults, the pITG has been shown to be involved in the visual processing of Arabic numbers (Conrad et al., 2020; Park et al., 2012; Shum et al., 2013; Skagenholt et al., 2018; Yeo et al., 2020), and is implicated in adult mathematics processing more generally (Amalric & Dehaene, 2016, 2019; Grotheer et al., 2018; Pinheiro-Chagas et al., 2018). Yet no clear number-specific responses have been found in the pITG in fMRI studies of children (Cantlon et al., 2011; Dehaene-Lambertz et al., 2018; Soltanlou et al., 2019). The absence of systematic activation of the pITG in children could suggest that these math-related neural responses develop later in childhood.

In adults, but not in children, the naturalistic mathematics task also yielded greater activation than the controlled mathematics task in the angular gyrus, that is often found in studies of mental calculation (Ansari, 2008; Arsalidou & Taylor, 2011; Dehaene et al., 2003; Delazer et al., 2005; Grabner et al., 2009; Ischebeck et al., 2006). The angular gyrus is generally seen as a semantic and conceptual hub, as it is involved in word and sentence comprehension, memory retrieval, spatial attention, and reasoning among other functions (e.g., Binder et al., 2009; Boylan et al., 2015; Price et al., 2015; Seghier, 2013; Zhou et al., 2018). Our finding provides converging evidence with previous studies that the angular gyrus has a protracted developmental trajectory in mathematics (Rivera et al., 2005; Soltanlou et al., 2018; Zamarian et al., 2009), just as observed in other domains such as reading (Booth et al., 2004; Church et al., 2008).

In children, within brain regions that are thought to respond to math semantics, the naturalistic and laboratory tasks differed in their neural signatures particularly in the anterior part of the right IPS. Many previous studies suggest that the right IPS matures faster and plays a greater role than the left IPS in young children’s early numerical development (Ansari, 2008; Cantlon et al., 2006; Matejko & Ansari, 2021; Vogel et al., 2015). Children appear to have a right lateralization bias for mathematics processing in the parietal cortex whereas activity in the left IPS emerges gradually over development and tracks the acquisition of formal symbolic numerical knowledge (Bugden et al., 2012; Cantlon et al., 2006; Emerson & Cantlon, 2015; Kaufmann et al., 2011). Our results indicate that naturalistic mathematics learning systematically engages a greater expanse of the dominant right IPS in children compared to the controlled math task, and mostly showed equivalent activation to the controlled task in the left IPS. One possibility is that children engage greater neural resources within the parietal mathematics network during the naturalistic task compared to the more stripped-down controlled task.

Naturalistic learning is a key phenomenon to understand in human brain development and, although this is a new area of enquiry, several studies have made progress toward comparing naturalistic learning with controlled laboratory performance in children (see Cantlon, 2020, p. 20; Vanderwal et al., 2019 for review). We found that a naturalistic mathematics lesson systematically evoked functional activation in brain regions that are classically involved in mathematics tasks. Our comparison of arithmetic versus grammar lessons yielded the predicted dissociation between math- and language-networks. The data revealed key functional similarities between math-related neural processes during qualitatively different tasks that probe the same arithmetic functions, one a naturalistic video lesson and the other a laboratory task. These findings suggest that regions of the intraparietal cortex process mathematical content when children are learning about mathematics in the real world. Moreover, the naturalistic math task engaged a broader expanse of the math network compared to the controlled math task, suggesting that naturalistic educational tasks may be more powerful than controlled tasks for eliciting activation within the math network – or for revealing functions that cut across math-specific and domain-general functions.

## Data availability statement

The data that support the findings of this study are openly available in OSF at https://osf.io/3a8hf/.

## Ethics approval statement

All participants gave informed consent after reading or being read consent information. The protocol was approved by the Carnegie Mellon Institutional Review Board.

## Appendices

### Appendix 1. MNI Peak Coordinates for the contrasts of the naturalistic math video lesson versus the controlled laboratory math task

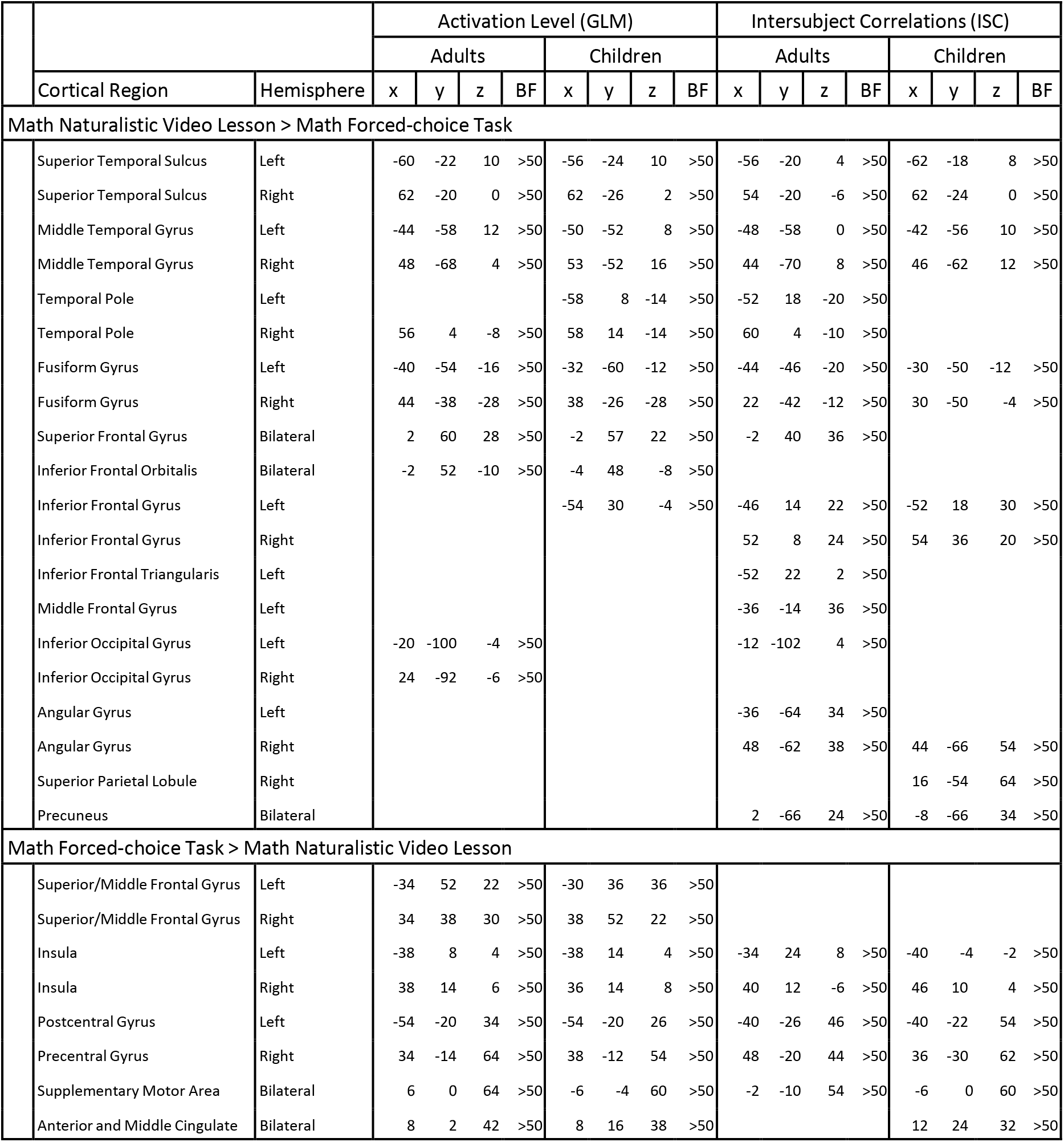

|  |  |  | Activation Level (GLM) |  |  |  |  |  |  |  | Intersubject Correlations (ISC) |  |  |  |  |  |  |  |
| --- | --- | --- | --- | --- | --- | --- | --- | --- | --- | --- | --- | --- | --- | --- | --- | --- | --- | --- |
|  |  |  | Adults |  |  |  | Children |  |  |  | Adults |  |  |  | Children |  |  |  |
|  | Cortical Region | Hemisphere | x | y | z | BF | x | y | z | BF | x | y | z | BF | x | y | z | BF |
| Math Naturalistic Video Lesson > Math Forced-choice Task |  |  |  |  |  |  |  |  |  |  |  |  |  |  |  |  |  |  |
|  | Superior Temporal Sulcus | Left | -60 | -22 | 10 | >50 | -56 | -24 | 10 | >50 | -56 | -20 | 4 | >50 | -62 | -18 | 8 | >50 |
|  | Superior Temporal Sulcus | Right | 62 | -20 | 0 | >50 | 62 | -26 | 2 | >50 | 54 | -20 | -6 | >50 | 62 | -24 | 0 | >50 |
|  | Middle Temporal Gyrus | Left | -44 | -58 | 12 | >50 | -50 | -52 | 8 | >50 | -48 | -58 | 0 | >50 | -42 | -56 | 10 | >50 |
|  | Middle Temporal Gyrus | Right | 48 | -68 | 4 | >50 | 53 | -52 | 16 | >50 | 44 | -70 | 8 | >50 | 46 | -62 | 12 | >50 |
|  | Temporal Pole | Left |  |  |  |  | -58 | 8 | -14 | >50 | -52 | 18 | -20 | >50 |  |  |  |  |
|  | Temporal Pole | Right | 56 | 4 | -8 | >50 | 58 | 14 | -14 | >50 | 60 | 4 | -10 | >50 |  |  |  |  |
|  | Fusiform Gyrus | Left | -40 | -54 | -16 | >50 | -32 | -60 | -12 | >50 | -44 | -46 | -20 | >50 | -30 | -50 | -12 | >50 |
|  | Fusiform Gyrus | Right | 44 | -38 | -28 | >50 | 38 | -26 | -28 | >50 | 22 | -42 | -12 | >50 | 30 | -50 | -4 | >50 |
|  | Superior Frontal Gyrus | Bilateral | 2 | 60 | 28 | >50 | -2 | 57 | 22 | >50 | -2 | 40 | 36 | >50 |  |  |  |  |
|  | Inferior Frontal Orbitalis | Bilateral | -2 | 52 | -10 | >50 | -4 | 48 | -8 | >50 |  |  |  |  |  |  |  |  |
|  | Inferior Frontal Gyrus | Left |  |  |  |  | -54 | 30 | -4 | >50 | -46 | 14 | 22 | >50 | -52 | 18 | 30 | >50 |
|  | Inferior Frontal Gyrus | Right |  |  |  |  |  |  |  |  | 52 | 8 | 24 | >50 | 54 | 36 | 20 | >50 |
|  | Inferior Frontal Triangularis | Left |  |  |  |  |  |  |  |  | -52 | 22 | 2 | >50 |  |  |  |  |
|  | Middle Frontal Gyrus | Left |  |  |  |  |  |  |  |  | -36 | -14 | 36 | >50 |  |  |  |  |
|  | Inferior Occipital Gyrus | Left | -20 | -100 | -4 | >50 |  |  |  |  | -12 | -102 | 4 | >50 |  |  |  |  |
|  | Inferior Occipital Gyrus | Right | 24 | -92 | -6 | >50 |  |  |  |  |  |  |  |  |  |  |  |  |
|  | Angular Gyrus | Left |  |  |  |  |  |  |  |  | -36 | -64 | 34 | >50 |  |  |  |  |
|  | Angular Gyrus | Right |  |  |  |  |  |  |  |  | 48 | -62 | 38 | >50 | 44 | -66 | 54 | >50 |
|  | Superior Parietal Lobule | Right |  |  |  |  |  |  |  |  |  |  |  |  | 16 | -54 | 64 | >50 |
|  | Precuneus | Bilateral |  |  |  |  |  |  |  |  | 2 | -66 | 24 | >50 | -8 | -66 | 34 | >50 |
| Math Forced-choice Task > Math Naturalistic Video Lesson |  |  |  |  |  |  |  |  |  |  |  |  |  |  |  |  |  |  |
|  | Superior/Middle Frontal Gyrus | Left | -34 | 52 | 22 | >50 | -30 | 36 | 36 | >50 |  |  |  |  |  |  |  |  |
|  | Superior/Middle Frontal Gyrus | Right | 34 | 38 | 30 | >50 | 38 | 52 | 22 | >50 |  |  |  |  |  |  |  |  |
|  | Insula | Left | -38 | 8 | 4 | >50 | -38 | 14 | 4 | >50 | -34 | 24 | 8 | >50 | -40 | -4 | -2 | >50 |
|  | Insula | Right | 38 | 14 | 6 | >50 | 36 | 14 | 8 | >50 | 40 | 12 | -6 | >50 | 46 | 10 | 4 | >50 |
|  | Postcentral Gyrus | Left | -54 | -20 | 34 | >50 | -54 | -20 | 26 | >50 | -40 | -26 | 46 | >50 | -40 | -22 | 54 | >50 |
|  | Precentral Gyrus | Right | 34 | -14 | 64 | >50 | 38 | -12 | 54 | >50 | 48 | -20 | 44 | >50 | 36 | -30 | 62 | >50 |
|  | Supplementary Motor Area | Bilateral | 6 | 0 | 64 | >50 | -6 | -4 | 60 | >50 | -2 | -10 | 54 | >50 | -6 | 0 | 60 | >50 |
|  | Anterior and Middle Cingulate | Bilateral | 8 | 2 | 42 | >50 | 8 | 16 | 38 | >50 |  |  |  |  | 12 | 24 | 32 | >50 |

### Appendix 2. MNI Peak Coordinates for the contrasts of the naturalistic math video lesson versus the naturalistic grammar video lesson

|  |  |  | Activation Level (GLM) |  |  |  |  |  |  |  | Intersubject Correlations (ISC) |  |  |  |  |  |  |  |
| --- | --- | --- | --- | --- | --- | --- | --- | --- | --- | --- | --- | --- | --- | --- | --- | --- | --- | --- |
|  |  |  | Adults |  |  |  | Children |  |  |  | Adults |  |  |  | Children |  |  |  |
|  | Cortical Region | Hemisphere | x | y | z | BF | x | y | z | BF | x | y | z | BF | x | y | z | BF |
| Math Naturalistic Video Lesson > Grammar Naturalistic Video Lesson |  |  |  |  |  |  |  |  |  |  |  |  |  |  |  |  |  |  |
|  | Superior Parietal Lobule | Left | -12 | -70 | 54 | >50 |  |  |  |  | -26 | -50 | 62 | >50 |  |  |  |  |
|  | Superior Parietal Lobule | Right | 54 | -28 | 58 | >50 |  |  |  |  | 28 | -52 | 62 | >50 |  |  |  |  |
|  | Inferior Parietal Lobule | Left | -46 | -40 | 56 | >50 | -36 | -46 | 42 | 14 | -54 | -28 | 44 | >50 | -32 | -40 | 48 | >50 |
|  | Inferior Parietal Lobule | Right | 46 | -46 | 54 | >50 | 38 | -42 | 46 | 49 |  |  |  |  | 38 | -40 | 52 | >50 |
|  | Inferior Temporal Gyrus | Left | -56 | -66 | -12 | >50 |  |  |  |  | -48 | -58 | -2 | >50 |  |  |  |  |
|  | Inferior Temporal Gyrus | Right |  |  |  |  | 56 | -60 | -18 | 26 | 56 | -58 | -14 | >50 |  |  |  |  |
|  | Left Sup Frontal Gyrus | Left | -28 | -4 | 66 | >50 |  |  |  |  | -22 | -8 | 52 | >50 |  |  |  |  |
|  | Right Inf Frontal Gyrus | Right | 54 | 14 | 42 | >50 |  |  |  |  | 58 | 14 | 26 | >50 | 52 | 4 | 30 | >50 |
|  | Calcarine/Lingual Gyrus | Right |  |  |  |  |  |  |  |  | 12 | -68 | 4 | >50 |  |  |  |  |
|  | Cuneus | Bilateral |  |  |  |  |  |  |  |  | 2 | -72 | 24 | >50 |  |  |  |  |
| Grammar Naturalistic Video Lesson > Math Naturalistic Video Lesson |  |  |  |  |  |  |  |  |  |  |  |  |  |  |  |  |  |  |
|  | Superior Temporal Sulcus | Left | -46 | -34 | 4 | >50 | -64 | -38 | 4 | >50 |  |  |  |  |  |  |  |  |
|  | Superior Temporal Sulcus | Right |  |  |  |  |  |  |  |  |  |  |  |  | 48 | -22 | -6 | >50 |
|  | Temporal Pole | Right | 58 | 2 | -12 | >50 | 48 | 2 | -14 | >50 |  |  |  |  | 34 | 12 | -42 | >50 |
|  | Temporo-Parietal Junction | Left | -46 | -58 | 16 | >50 | -42 | -56 | 28 | 49 |  |  |  |  | -42 | -60 | 20 | >50 |
|  | Temporo-Parietal Junction | Right | 44 | -52 | 24 | >50 | 50 | -66 | 20 | 25 |  |  |  |  | 50 | -48 | 24 | >50 |
|  | Inferior Occipital Gyrus | Left | -20 | -96 | -10 | >50 |  |  |  |  |  |  |  |  |  |  |  |  |
|  | Inferior Occipital Gyrus | Right | 18 | -92 | -4 | >50 |  |  |  |  |  |  |  |  |  |  |  |  |
|  | Insula | Left |  |  |  |  | -38 | -20 | 14 | >50 | -36 | 14 | -8 | >50 | -38 | 14 | 0 | >50 |
|  | Inferior Frontal Opercularis | Left | -36 | 14 | 22 | >50 |  |  |  |  |  |  |  |  |  |  |  |  |
|  | Inferior Frontal Opercularis | Right |  |  |  |  |  |  |  |  |  |  |  |  | 50 | 14 | 10 | >50 |
|  | Inferior Frontal Triangularis | Left |  |  |  |  |  |  |  |  | -44 | 36 | 2 | >50 |  |  |  |  |
|  | Superior Frontal Gyrus | Left |  |  |  |  |  |  |  |  | -24 | 32 | 42 | >50 |  |  |  |  |
|  | Superior Frontal Gyrus | Right |  |  |  |  |  |  |  |  | 18 | 40 | 34 | >50 |  |  |  |  |
|  | Middle Frontal Gyrus | Left |  |  |  |  |  |  |  |  |  |  |  |  | -30 | 48 | 8 | >50 |
|  | Middle Frontal Gyrus | Right |  |  |  |  |  |  |  |  |  |  |  |  | 34 | 40 | 8 | >50 |
|  | Left Postcentral Gyrus | Left |  |  |  |  |  |  |  |  | -54 | -16 | 46 | >50 | -40 | -22 | 52 | >50 |

## References

Amalric, M. (2021). Cerebral underpinning of advanced mathematical activity. In Heterogeneous Contributions to Numerical Cognition (pp. 71–92). Elsevier. https://doi.org/10.1016/B978-0-12-817414-2.00009-9

Amalric, M., & Dehaene, S. (2016). Origins of the brain networks for advanced mathematics in expert mathematicians. Proceedings of the National Academy of Sciences, 201603205. https://doi.org/10.1073/pnas.1603205113

Amalric, M., & Dehaene, S. (2018). Cortical circuits for mathematical knowledge: Evidence for a major subdivision within the brain’s semantic networks. Phil. Trans. R. Soc. B, 373(1740), 20160515. https://doi.org/10.1098/rstb.2016.0515

Amalric, M., & Dehaene, S. (2019). A distinct cortical network for mathematical knowledge in the human brain. NeuroImage, 189, 19–31. https://doi.org/10.1016/j.neuroimage.2019.01.001

Amalric, M., Roveyaz, P., & Dehaene, S. (in prep). Can a short math video enhance the brain’s mathematical networks?

Ansari, D. (2008). Effects of development and enculturation on number representation in the brain. Nature Reviews Neuroscience, 9(4), 278–291. https://doi.org/10.1038/nrn2334

Ansari, D. (2016). The neural roots of mathematical expertise. Proceedings of the National Academy of Sciences, 113(18), 4887–4889. https://doi.org/10.1073/pnas.1604758113

Arsalidou, M., Pawliw-Levac, M., Sadeghi, M., & Pascual-Leone, J. (2018). Brain areas associated with numbers and calculations in children: Meta-analyses of fMRI studies. Developmental Cognitive Neuroscience, 30, 239–250. https://doi.org/10.1016/j.dcn.2017.08.002

Arsalidou, M., & Taylor, M. J. (2011). Is 2+2=4? Meta-analyses of brain areas needed for numbers and calculations. NeuroImage, 54(3), 2382–2393. https://doi.org/10.1016/j.neuroimage.2010.10.009

Ashkenazi, S., Rosenberg-Lee, M., Tenison, C., & Menon, V. (2012). Weak task-related modulation and stimulus representations during arithmetic problem solving in children with developmental dyscalculia. Developmental Cognitive Neuroscience, 2, S152–S166. https://doi.org/10.1016/j.dcn.2011.09.006

Baldo, J. V., & Dronkers, N. F. (2007). Neural correlates of arithmetic and language comprehension: A common substrate? Neuropsychologia, 45(2), 229–235. https://doi.org/10.1016/j.neuropsychologia.2006.07.014

Binder, J. R., Desai, R. H., Graves, W. W., & Conant, L. L. (2009). Where Is the Semantic System? A Critical Review and Meta-Analysis of 120 Functional Neuroimaging Studies. Cerebral Cortex, 19(12), 2767–2796. https://doi.org/10.1093/cercor/bhp055

Booth, J. R., Burman, D. D., Meyer, J. R., Gitelman, D. R., Parrish, T. B., & Mesulam, M. M. (2004). Development of Brain Mechanisms for Processing Orthographic and Phonologic Representations. Journal of Cognitive Neuroscience, 16(7), 1234–1249. https://doi.org/10.1162/0898929041920496

Boylan, C., Trueswell, J. C., & Thompson-Schill, S. L. (2015). Compositionality and the angular gyrus: A multi-voxel similarity analysis of the semantic composition of nouns and verbs. Neuropsychologia, 78, 130–141. https://doi.org/10.1016/j.neuropsychologia.2015.10.007

Bugden, S., Price, G. R., McLean, D. A., & Ansari, D. (2012). The role of the left intraparietal sulcus in the relationship between symbolic number processing and children’s arithmetic competence. Developmental Cognitive Neuroscience, 2(4), 448–457. https://doi.org/10.1016/j.dcn.2012.04.001

Cantlon, J. F. (2015). Chapter 9—Analog Origins of Numerical Concepts. In D. C. Geary, D. B. Berch, & K.M. Koepke (Eds.), Mathematical Cognition and Learning (Vol. 1, pp. 225–251). Elsevier. https://doi.org/10.1016/B978-0-12-420133-0.00009-0

Cantlon, J. F. (2020). The balance of rigor and reality in developmental neuroscience. NeuroImage, 216, 116464. https://doi.org/10.1016/j.neuroimage.2019.116464

Cantlon, J. F., Brannon, E. M., Carter, E. J., & Pelphrey, K. A. (2006). Functional Imaging of Numerical Processing in Adults and 4-y-Old Children. PLOS Biology, 4(5), e125. https://doi.org/10.1371/journal.pbio.0040125

Cantlon, J. F., & Li, R. (2013). Neural Activity during Natural Viewing of Sesame Street Statistically Predicts Test Scores in Early Childhood. PLoS Biology, 11(1), e1001462. https://doi.org/10.1371/journal.pbio.1001462

Cantlon, J. F., Pinel, P., Dehaene, S., & Pelphrey, K. A. (2011). Cortical Representations of Symbols, Objects, and Faces Are Pruned Back during Early Childhood. Cerebral Cortex, 21(1), 191–199. https://doi.org/10.1093/cercor/bhq078

Church, J. A., Coalson, R. S., Lugar, H. M., Petersen, S. E., & Schlaggar, B. L. (2008). A Developmental fMRI Study of Reading and Repetition Reveals Changes in Phonological and Visual Mechanisms Over Age. Cerebral Cortex, 18(9), 2054–2065. https://doi.org/10.1093/cercor/bhm228

Conrad, B. N., Wilkey, E. D., Yeo, D. J., & Price, G. R. (2020). Network topology of symbolic and nonsymbolic number comparison. Network Neuroscience, 4(3), 714–745. https://doi.org/10.1162/netn_a_00144

Dastjerdi, M., Ozker, M., Foster, B. L., Rangarajan, V., & Parvizi, J. (2013). Numerical processing in the human parietal cortex during experimental and natural conditions. Nature Communications, 4, ncomms3528. https://doi.org/10.1038/ncomms3528

De Smedt, B., Noël, M.-P., Gilmore, C., & Ansari, D. (2013). How do symbolic and non-symbolic numerical magnitude processing skills relate to individual differences in children’s mathematical skills? A review of evidence from brain and behavior. Trends in Neuroscience and Education, 2(2), 48–55. https://doi.org/10.1016/j.tine.2013.06.001

Dehaene, S. (2004). Arithmetic and the brain. Current Opinion in Neurobiology, 14(2), 218–224. https://doi.org/10.1016/j.conb.2004.03.008

Dehaene, S., & Cohen, L. (2007). Cultural Recycling of Cortical Maps. Neuron, 56(2), 384–398. https://doi.org/10.1016/j.neuron.2007.10.004

Dehaene, S., Piazza, M., Pinel, P., & Cohen, L. (2003). THREE PARIETAL CIRCUITS FOR NUMBER PROCESSING. Cognitive Neuropsychology, 20(3–6), 487–506. https://doi.org/10.1080/02643290244000239

Dehaene-Lambertz, G., Monzalvo, K., & Dehaene, S. (2018). The emergence of the visual word form:Longitudinal evolution of category-specific ventral visual areas during reading acquisition. PLOS Biology, 16(3), e2004103. https://doi.org/10.1371/journal.pbio.2004103

Delazer, M., Domahs, F., Bartha, L., Brenneis, C., Lochy, A., Trieb, T., & Benke, T. (2003). Learning complex arithmetic—An fMRI study. Cognitive Brain Research, 18(1), 76–88. https://doi.org/10.1016/j.cogbrainres.2003.09.005

Delazer, M., Ischebeck, A., Domahs, F., Zamarian, L., Koppelstaetter, F., Siedentopf, C. M., Kaufmann, L., Benke, T., & Felber, S. (2005). Learning by strategies and learning by drill—Evidence from an fMRI study. NeuroImage, 25(3), 838–849. https://doi.org/10.1016/j.neuroimage.2004.12.009

Dienes, Z. (2014). Using Bayes to get the most out of non-significant results. Frontiers in Psychology, 5. https://doi.org/10.3389/fpsyg.2014.00781

Emerson, R. W., & Cantlon, J. F. (2015). Continuity and change in children’s longitudinal neural responses to numbers. Developmental Science, 18(2), 314–326. https://doi.org/10.1111/desc.12215

Grabner, R. H., Ansari, D., Koschutnig, K., Reishofer, G., Ebner, F., & Neuper, C. (2009). To retrieve or to calculate? Left angular gyrus mediates the retrieval of arithmetic facts during problem solving. Neuropsychologia, 47(2), 604–608. https://doi.org/10.1016/j.neuropsychologia.2008.10.013

Grotheer, M., Jeska, B., & Grill-Spector, K. (2018). A preference for mathematical processing outweighs the selectivity for Arabic numbers in the inferior temporal gyrus. NeuroImage, 175, 188–200. https://doi.org/10.1016/j.neuroimage.2018.03.064

Hawes, Z., Sokolowski, H. M., Ononye, C. B., & Ansari, D. (2019). Neural underpinnings of numerical and spatial cognition: An fMRI meta-analysis of brain regions associated with symbolic number, arithmetic, and mental rotation. Neuroscience & Biobehavioral Reviews, 103, 316–336. https://doi.org/10.1016/j.neubiorev.2019.05.007

Holloway, I. D., Price, G. R., & Ansari, D. (2010). Common and segregated neural pathways for the processing of symbolic and nonsymbolic numerical magnitude: An fMRI study. NeuroImage, 49(1), 1006–1017. https://doi.org/10.1016/j.neuroimage.2009.07.071

Houdé, O., Rossi, S., Lubin, A., & Joliot, M. (2010). Mapping numerical processing, reading, and executive functions in the developing brain: An fMRI meta-analysis of 52 studies including 842 children. Developmental Science, 13(6), 876–885. https://doi.org/10.1111/j.1467-7687.2009.00938.x

Ischebeck, A., Zamarian, L., Siedentopf, C., Koppelstätter, F., Benke, T., Felber, S., & Delazer, M. (2006). How specifically do we learn? Imaging the learning of multiplication and subtraction. NeuroImage, 30(4), 1365–1375. https://doi.org/10.1016/j.neuroimage.2005.11.016

Jeffreys, H. (1961). The Theory of Probability (3rd ed). Oxford University Press.

Jeon, H.-A., Kuhl, U., & Friederici, A. D. (2019). Mathematical expertise modulates the architecture of dorsal and cortico-thalamic white matter tracts. Scientific Reports, 9(1), 1–11. https://doi.org/10.1038/s41598-019-43400-6

Jolles, D., Supekar, K., Richardson, J., Tenison, C., Ashkenazi, S., Rosenberg-Lee, M., Fuchs, L., & Menon, V. (2016). Reconfiguration of parietal circuits with cognitive tutoring in elementary school children. Cortex, 83, 231–245. https://doi.org/10.1016/j.cortex.2016.08.004

Kanjlia, S., Lane, C., Feigenson, L., & Bedny, M. (2016). Absence of visual experience modifies the neural basis of numerical thinking. Proceedings of the National Academy of Sciences, 201524982. https://doi.org/10.1073/pnas.1524982113

Kaufmann, L., Wood, G., Rubinsten, O., & Henik, A. (2011). Meta-Analyses of Developmental fMRI Studies Investigating Typical and Atypical Trajectories of Number Processing and Calculation. Developmental Neuropsychology, 36(6), 763–787. https://doi.org/10.1080/87565641.2010.549884

Kersey, A. J., Wakim, K.-M., Li, R., & Cantlon, J. F. (2019). Developing, mature, and unique functions of the child’s brain in reading and mathematics. Developmental Cognitive Neuroscience, 39, 100684. https://doi.org/10.1016/j.dcn.2019.100684

Klessinger, N., Szczerbinski, M., & Varley, R. (2007). Algebra in a man with severe aphasia. Neuropsychologia, 45(8), 1642–1648. https://doi.org/10.1016/j.neuropsychologia.2007.01.005

Lerner, Y., Scherf, K. S., Katkov, M., Hasson, U., & Behrmann, M. (2021). Changes in Cortical Coherence Supporting Complex Visual and Social Processing in Adolescence. Journal of Cognitive Neuroscience, 33(11), 2215–2230. https://doi.org/10.1162/jocn_a_01756

Maruyama, M., Pallier, C., Jobert, A., Sigman, M., & Dehaene, S. (2012). The cortical representation of simple mathematical expressions. NeuroImage, 61(4), 1444–1460. https://doi.org/10.1016/j.neuroimage.2012.04.020

Matejko, A. A., & Ansari, D. (2021). Shared Neural Circuits for Visuospatial Working Memory and Arithmetic in Children and Adults. Journal of Cognitive Neuroscience, 1–17. https://doi.org/10.1162/jocn_a_01695

Monti, M. M., Parsons, L. M., & Osherson, D. N. (2012). Thought Beyond Language: Neural Dissociation of Algebra and Natural Language. Psychological Science, 23(8), 914–922. https://doi.org/10.1177/0956797612437427

Nastase, S. A., Gazzola, V., Hasson, U., & Keysers, C. (2019). Measuring shared responses across subjects using intersubject correlation. Social Cognitive and Affective Neuroscience, 14(6), 667–685. https://doi.org/10.1093/scan/nsz037

O’Boyle, M. W., Cunnington, R., Silk, T. J., Vaughan, D., Jackson, G., Syngeniotis, A., & Egan, G. F. (2005). Mathematically gifted male adolescents activate a unique brain network during mental rotation. Cognitive Brain Research, 25(2), 583–587. https://doi.org/10.1016/j.cogbrainres.2005.08.004

Park, J., Hebrank, A., Polk, T. A., & Park, D. C. (2012). Neural dissociation of number from letter recognition and its relationship to parietal numerical processing. Journal of Cognitive Neuroscience, 24(1), 39–50.

Peters, L., & De Smedt, B. (2018). Arithmetic in the developing brain: A review of brain imaging studies. Developmental Cognitive Neuroscience, 30, 265–279. https://doi.org/10.1016/j.dcn.2017.05.002

Pinel, P., Thirion, B., Meriaux, S., Jobert, A., Serres, J., Le Bihan, D., Poline, J.-B., & Dehaene, S. (2007). Fast reproducible identification and large-scale databasing of individual functional cognitive networks. BMC Neuroscience, 8(1), 91. https://doi.org/10.1186/1471-2202-8-91

Pinheiro-Chagas, P., Daitch, A., Parvizi, J., & Dehaene, S. (2018). Brain Mechanisms of Arithmetic: A Crucial Role for Ventral Temporal Cortex. Journal of Cognitive Neuroscience, 30(12), 1757–1772. https://doi.org/10.1162/jocn_a_01319

Prescott, J., Gavrilescu, M., Cunnington, R., O’Boyle, M. W., & Egan, G. F. (2010). Enhanced brain connectivity in math-gifted adolescents: An fMRI study using mental rotation. Cognitive Neuroscience, 1(4), 277–288. https://doi.org/10.1080/17588928.2010.506951

Price, A. R., Bonner, M. F., Peelle, J. E., & Grossman, M. (2015). Converging Evidence for the Neuroanatomic Basis of Combinatorial Semantics in the Angular Gyrus. Journal of Neuroscience, 35(7), 3276–3284. https://doi.org/10.1523/JNEUROSCI.3446-14.2015

Price, G. R., Holloway, I., Räsänen, P., Vesterinen, M., & Ansari, D. (2007). Impaired parietal magnitude processing in developmental dyscalculia. Current Biology, 17(24), R1042–R1043. https://doi.org/10.1016/j.cub.2007.10.013

Price, G. R., Mazzocco, M. M. M., & Ansari, D. (2013). Why Mental Arithmetic Counts: Brain Activation during Single Digit Arithmetic Predicts High School Math Scores. Journal of Neuroscience, 33(1), 156–163. https://doi.org/10.1523/JNEUROSCI.2936-12.2013

Richardson, H., Lisandrelli, G., Riobueno-Naylor, A., & Saxe, R. (2018). Development of the social brain from age three to twelve years. Nature Communications, 9(1), 1027. https://doi.org/10.1038/s41467-018-03399-2

Rivera, S. M., Reiss, A. L., Eckert, M. A., & Menon, V. (2005). Developmental Changes in Mental Arithmetic: Evidence for Increased Functional Specialization in the Left Inferior Parietal Cortex. Cerebral Cortex, 15(11), 1779–1790. https://doi.org/10.1093/cercor/bhi055

Rouder, J. N., Speckman, P. L., Sun, D., Morey, R. D., & Iverson, G. (2009). Bayesian t tests for accepting and rejecting the null hypothesis. Psychonomic Bulletin & Review, 16(2), 225–237. https://doi.org/10.3758/PBR.16.2.225

Schönbrodt, F.D., Wagenmakers, E.-J., Zehetleitner, M., Perugini, M., & Psychologische Methodenleer (Psychologie, FMG). (2017). Sequential Hypothesis Testing With Bayes Factors: Efficiently Testing Mean Differences. Psychological Methods, 22(2), 322–339. https://doi.org/10.1037/met0000061

Seghier, M. L. (2013). The Angular Gyrus: Multiple Functions and Multiple Subdivisions. The Neuroscientist, 19(1), 43–61. https://doi.org/10.1177/1073858412440596

Shum, J., Hermes, D., Foster, B. L., Dastjerdi, M., Rangarajan, V., Winawer, J., Miller, K. J., & J. Parvizi. (2013). A Brain Area for Visual Numerals. Journal of Neuroscience, 33(16), 6709–6715. https://doi.org/10.1523/JNEUROSCI.4558-12.2013

Simon, O., Mangin, J.-F., Cohen, L., Le Bihan, D., & Dehaene, S. (2002). Topographical layout of hand, eye, calculation, and language-related areas in the human parietal lobe. Neuron, 33(3), 475–487.

Skagenholt, M., Träff, U., Västfjäll, D., & Skagerlund, K. (2018). Examining the Triple Code Model in numerical cognition: An fMRI study. PLOS ONE, 13(6), e0199247. https://doi.org/10.1371/journal.pone.0199247

Soltanlou, M., Coldea, A., Artemenko, C., Ehlis, A.-C., Fallgatter, A. J., Nuerk, H.-C., & Dresler, T. (2019). No Difference in the Neural Underpinnings of Number and Letter Copying in Children: Bayesian Analysis of Functional Near-Infrared Spectroscopy Data. Mind, Brain, and Education, 13(4), 313–325. https://doi.org/10.1111/mbe.12225

Soltanlou, M., Sitnikova, M. A., Nuerk, H.-C., & Dresler, T. (2018). Applications of Functional Near-Infrared Spectroscopy (fNIRS) in Studying Cognitive Development: The Case of Mathematics and Language. Frontiers in Psychology, 9, 277. https://doi.org/10.3389/fpsyg.2018.00277

Uddin, L. Q., & Menon, V. (2009). The anterior insula in autism: Under-connected and under-examined. Neuroscience & Biobehavioral Reviews, 33(8), 1198–1203. https://doi.org/10.1016/j.neubiorev.2009.06.002

Vanderwal, T., Eilbott, J., & Castellanos, F. X. (2019). Movies in the magnet: Naturalistic paradigms in developmental functional neuroimaging. Developmental Cognitive Neuroscience, 36, 100600. https://doi.org/10.1016/j.dcn.2018.10.004

Vogel, S. E., Goffin, C., & Ansari, D. (2015). Developmental specialization of the left parietal cortex for the semantic representation of Arabic numerals: An fMR-adaptation study. Developmental Cognitive Neuroscience, 12, 61–73. https://doi.org/10.1016/j.dcn.2014.12.001

Williams, M. N., Bååth, R. A., & Philipp, M. C. (2017). Using Bayes Factors to Test Hypotheses in Developmental Research. Research in Human Development, 14(4), 321–337. https://doi.org/10.1080/15427609.2017.1370964

Yeo, D. J., Pollack, C., Merkley, R., Ansari, D., & Price, G. R. (2020). The “Inferior Temporal Numeral Area” distinguishes numerals from other character categories during passive viewing: A representational similarity analysis. NeuroImage, 214, 116716. https://doi.org/10.1016/j.neuroimage.2020.116716

Yeo, D. J., Wilkey, E. D., & Price, G. R. (2017). The search for the number form area: A functional neuroimaging meta-analysis. Neuroscience & Biobehavioral Reviews. https://doi.org/10.1016/j.neubiorev.2017.04.027

Zamarian, L., Ischebeck, A., & Delazer, M. (2009). Neuroscience of learning arithmetic—Evidence from brain imaging studies. Neuroscience & Biobehavioral Reviews, 33(6), 909–925. https://doi.org/10.1016/j.neubiorev.2009.03.005

Zhang, H., Chen, C., & Zhou, X. (2012). Neural correlates of numbers and mathematical terms. NeuroImage, 60(1), 230–240. https://doi.org/10.1016/j.neuroimage.2011.12.006

Zhou, X., Li, M., Li, L., Zhang, Y., Cui, J., Liu, J., & Chen, C. (2018). The semantic system is involved in mathematical problem solving. NeuroImage, 166, 360–370. https://doi.org/10.1016/j.neuroimage.2017.11.017

